# ONC201 in combination with ATR inhibitor ceralasertib exhibits potent synergy in ovarian cancer cells

**DOI:** 10.64898/2026.09.23.753805

**Authors:** Maryam Ghandali, Kadir Mert Selvi, Kelsey E. Huntington, Andrew George, Anna Ochsner, Benedito A. Carneiro, Don S. Dizon, Leiqing Zhang, Wafik S. El-Deiry

## Abstract

Ovarian cancer stands out as one of the most lethal gynecological cancers among women in the United States. While patients with ovarian cancer often show positive responses to treatment, it is common for the cancer to reappear and develop resistance to treatment, so it is important to explore and create novel combinations to increase patient survival rates. ATRi’s (Ataxia telangiectasia and Rad3-related inhibitors) and ONC201 monotherapies have shown efficacy in ovarian cancer in preclinical models. We hypothesized that combining ATRi’s with ONC201 would enhance this efficacy. ONC201 induces apoptosis through the TRAIL-pathway following activation of the integrated stress response and inactivation of Akt which is upregulated in ovarian cancer. Six human ovarian cancer cell lines were treated alone and with the novel drug combination of ceralasertib (ATRi) and ONC201. We found an IC50 range of 0.88-20.16 μM for ONC201 and IC50 range of 0.55-54.75 μM for ceralasertib. The combination of ONC201 plus ceralasertib resulted in PARP cleavage consistent with apoptosis along with reduction in Bcl-XL, Xiap and p-Akt expression after 72 hours of treatment. Cytokine profiling showed a decrease in markers of inflammation and tumor progression. Our data provide evidence of synergy from the combination of ATRi ceralasertib and ONC201 against Ovarian cancer cells that merits further investigation including clinical translation.

## Introduction

Ovarian cancer stands is a particularly lethal gynecological cancer, often diagnosed at an advanced stage, among women in the United States. In 2024, there were estimated to be 19,680 new cases and 12,740 deaths from ovarian cancer in the US (1). Roughly 70% of individuals with ovarian cancer experience a recurrence following their initial treatment (2). Therefore, it is important to find new treatment combinations to enhance the long-term results and prolong patient survival.

ONC201 (TIC10) is the founding member of a novel class of small molecule anti-cancer drugs known as imipridones. Imipridones stimulate the integrated stress response, specifically by engaging the eIF2α/ATF4 pathway. Additionally, they induce Death Receptor 5 and its ligand the tumor necrosis factor (TNF)-related apoptosis-inducing ligand (TRAIL) thereby orchestrating a cascade of tumor regression and cell death across a diverse array of cancer models (3).

ONC201 functions as a compound that can simultaneously inhibit both MAPK/ERK signaling and Akt pathways (4). The PI3K/AKT pathway has an important role in oncogenic signaling, and there is substantial evidence showing that these pathways are often disrupted in numerous human malignancies (5).

Consequently, activation of Akt leads to activation or inhibition of a considerable number of downstream molecules that, in turn, contribute to cancer progression and resistance to anti-cancer treatments in addition to facilitating cancer metastasis (6). Roughly 70% of ovarian cancers have abnormally upregulated PI3K/AKT/mTOR signaling (7).

ONC201 has been investigated in a range of different preclinical and clinical settings covering a wide array of hematological and solid cancers. This includes studies on acute myeloid leukemia, brain cancer, colorectal cancer, breast cancer, and ovarian cancer (8). It has been shown that ONC201 is effective in inducing cancer cell death in both low-grade and high-grade ovarian cancer cell models (9). This makes ONC201, as a possible Akt inhibitory molecule, an appealing treatment option to test in ovarian malignancies.

ATR is a serine/threonine kinase that plays an essential role in the DNA damage response (DDR) by directing downstream cellular signaling following DNA damage resulting in single-stranded DNA structures (10,11). Inhibiting ATR kinase has potential to target and selectively kill genomically unstable cancer cells that are already under replication stress (12,13).

Therefore, various ATR inhibitors (ATRi) are actively being tested in multiple preclinical and clinical trials (14). ATRi’s have also shown promising results when tested on ovarian cancer (15,16).

AZD6738 (ceralasertib) is an effective and orally bioavailable ATRi that is in numerous clinical trials as a single agent or in combination therapies for multiple cancer types (17). Evidence indicates that signaling pathways with an “oncogenic” role in cell cycle regulation, such as the PI3K/AKT and DDR pathways, regulate and activate one another (18). In response to DNA damage, sensor proteins involved in DDR like ATM, ATR, and PARP are found to be engaged with the activation of AKT, and PI3K/AKT signaling which may contribute to resistance to DDR therapies (19). Thus, employing drugs that target both the PI3K/AKT and DDR pathways in combination may offer a potentially useful method for overcoming resistance and improving therapeutic efficacy in ovarian cancer (18,19). It has been shown that ONC201 and its derivative drugs work synergistically with some DDR inhibitors (20).

To further develop new combination strategies for the treatment of ovarian cancer, we hypothesized that the combination of ATRi ceralasertib and imipiridone ONC201 may be an effective combination for the treatment of ovarian cancer in preclinical models.

## Materials and methods

### Cell Lines

The TOV21G, OVCAR3, A2780, CAOV3, and SKOV3 cell lines were purchased from American Type Culture Collection (ATCC, Manassas, VA). The KURAMOCHI cell line was purchased from JCRB (JCRB0098). The TOV21G, OVCAR3, A2780, and KURAMOCHI cell lines were cultured in RPMI 1640 Medium, 10% fetal bovine serum, and 1% penicillin-streptomycin were added to this medium. The CAOV3 and SKOV3 cell lines were cultured in DMEM medium with 10% fetal bovine serum and 1% penicillin-streptomycin. Cells were incubated at 37°C in a 5% CO_2_ incubator.

### Cell viability and synergy assessment

A total of 5×10^3^ cells were seeded in 96 well plates in appropriate culture medium as mentioned above. Plates were kept overnight in 5% CO_2_ at 37°C. Cells were treated with selected drugs the next day. Cell viability was examined by Cell Titer-Glo (CTG, Promega, Madison, WI) after 72 hours of treatment. Graph pad prism version 9 (San Diego, CA, USA) was used to calculate Half maximal inhibitory concentration (IC50). Synergy was determined using Compusyn software (ComboSyn,Inc.).

### Immunoblotting

A total of 5×10^5^ cells-1×10^6^ were seeded in each well of a 6 well plate and kept overnight at 37℃ with 5% CO_2_. The following day cells were treated with desired drugs or vehicles and were incubated at 37°C. After the desired amount of time, cells were detached by using Protein lysis buffer. BCA protein assay kit (Thermo Fisher Scientific, Carlsbad, CA, USA) was used to calculate protein concentration. The same amount of protein was loaded and run in NuPAGE 4-12% Bis-Tris Gel (Thermo Fisher Scientific) and then proteins were transferred to PVDF membranes. 5% non-fat milk was used to block membranes. The membranes were kept overnight in primary antibodies that were diluted using 5% non-fat milk at 4℃. The next day secondary antibodies diluted in 5% non-fat milk were added to washed membranes. The antibodies used for western blot analysis were cPARP, Bcl-XL, Phospho-Akt (Ser473), XIAP, cIAP-1, DR5, vinculin all obtained from cell signaling. Anti-ATF-4 was obtained from Proteintech. CLPX antibody was obtained from NSJ bioreagents, Ran antibody from BD Biosciences and β-Actin was purchased from Sigma-Aldrich. Anti-Mouse and Anti-Rabbit were used as secondary antibodies.

| Antibody | Catalog number | Company | Dilution |
| --- | --- | --- | --- |
| Cleaved PARP (Asp214) (19F4) | 9546 | Cell Signaling | 1:1000 |
| Phospho-Akt (Ser473) (D9E) | 4060 | Cell Signaling | 1:1000 |
| XIAP | 2042 | Cell Signaling | 1:1000 |
| c-IAP1 (D5G9) | 7065 | Cell signaling | 1:1000 |
| DR5 (D4E9) | 8074 | Cell signaling | 1:1000 |
| Vinculin E1E9V | 13901 | Cell signaling | 1:1000 |
| Bcl-xL (54H6) | 2764 | Cell signaling | 1:1000 |
| ATF-4 | 10835-1-AP | Protein Tech | 1:1000 |
| ClpX | RQ5618 | NSJ bio reagents | 1:1000 |
| Ran | 610341 | BD Transduction Laboratories | 1:5000 |
| B-actin | A5441 | Sigma Aldrich | 1:5000 |

### Cytokine profiling

24 well plates were seeded with 1×10^5^ cells and incubated overnight with 5% CO_2_ at 37℃. Cells were treated with desired drugs, and after 24 or 48 hours of treatment, the supernatant was collected and used in luminex 200 for cytokine profiling. On the Luminex 200 instrument, a Human Premixed Multi-Analyte kit (R&D Systems, Inc., Minneapolis, MN) was used and the reported cytokine concentrations were in pg/ml. Cytokine concentrations given by the Luminex 200 as below the detection threshold were divided by 10 and this was used as the lower limit for comparison with other treatment conditions with the same cytokine. Cytokines above the upper limit of detection were used as the number given by the Luminex instrument. Cytokines without any detectable value were removed from analysis. Graph pad prism version 9 (San Diego, CA, USA) was used for statistical analysis of data.

## Results

### Reduced cell viability after single treatment with ONC201 and ceralasertib in ovarian cancer cell lines

To determine the killing effect of individual drugs in ovarian cancer cell lines, 6 cell lines were treated with single-agent ONC201 and ceralasertib. Cells were incubated for 72 hours, then analyzed with CTG. Half maximal inhibitory concentration (IC50) was determined for each drug in each cell line. The results including dose-response curves based on viability are shown in **Figure 1**. Loss of ovarian tumor cell viability was observed with both ONC201 and Ceralasertib. The IC50 values and known histologic type/p53 status of each cell line is listed in **Table 1** (22,23). We observed an IC50 range of 0.88-20.16 μM for ONC201 and 0.55-54.75 μM for Ceralasertib among the tested ovarian tumor cell lines.

**Figure 1.**
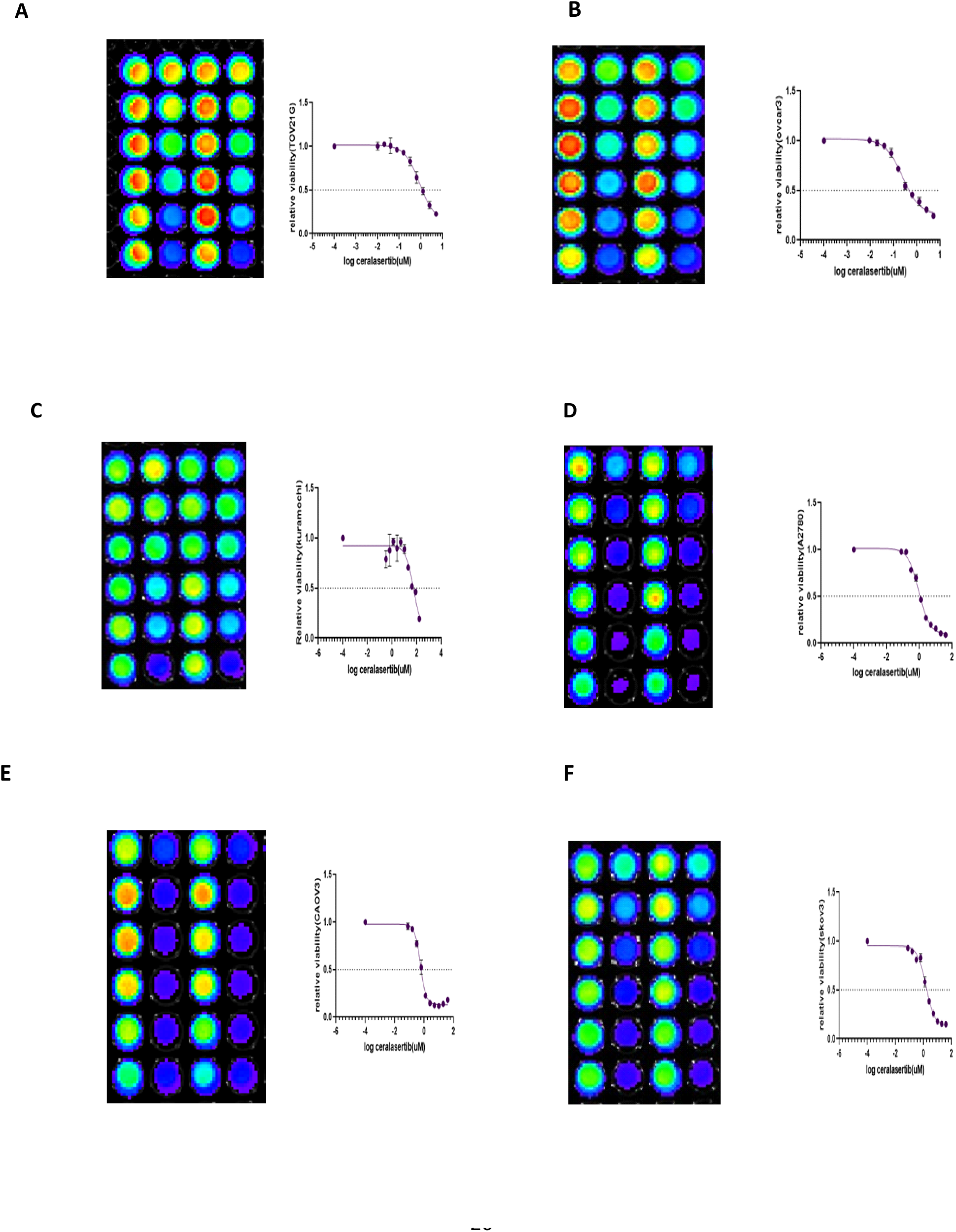
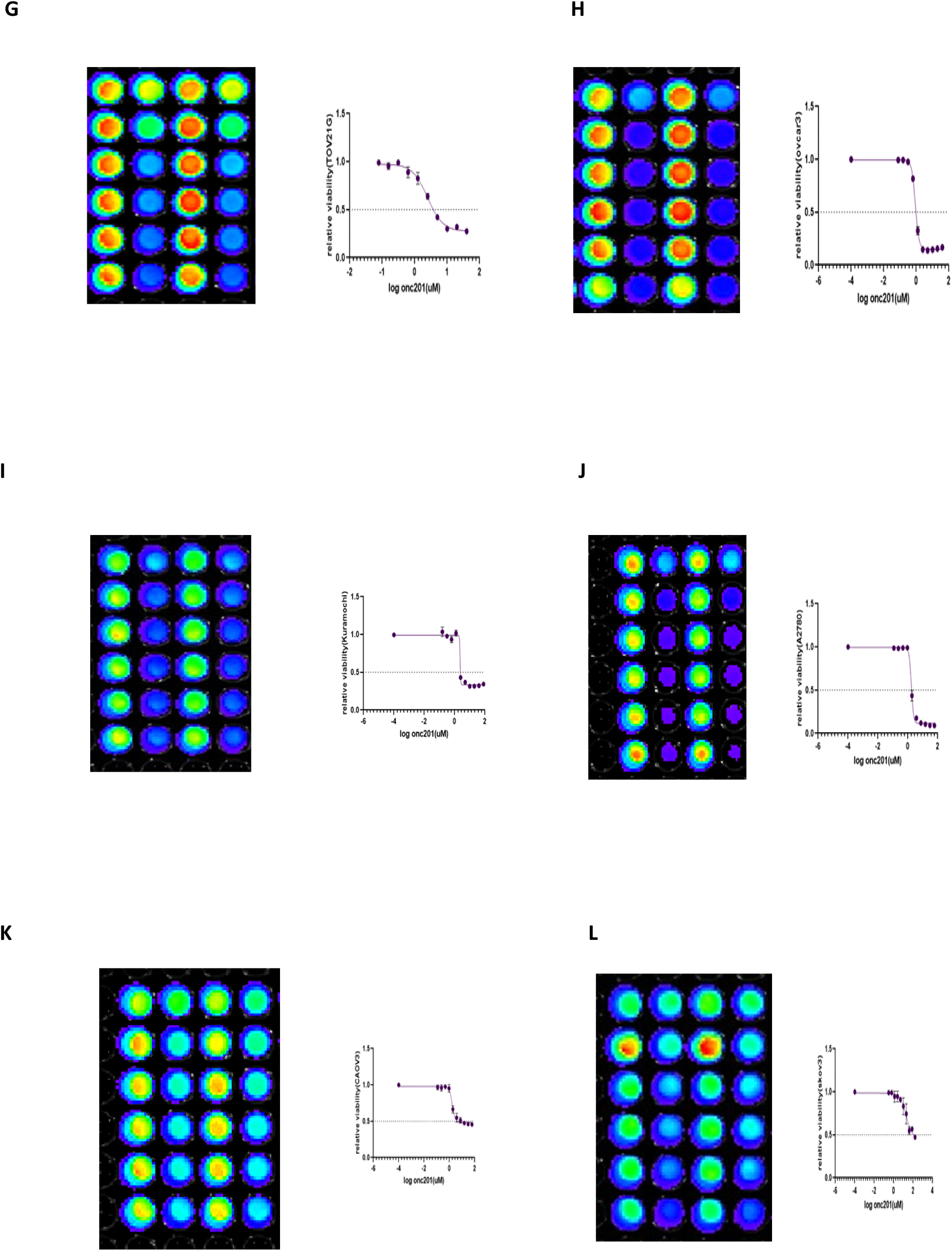
Ceralasertib and ONC201 reduce the viability of ovarian cancer cell lines. TOV21G, OVCAR3, Kuramochi, A2780, CAOV3 and skov3 cell line were cultured with increasing dosage of each drug, cell titer Glo viability assay were performed after 72 hours of treatment as described in Methods (each cell titer GLO cell viability assay pictures show the duplicate increasing dosage of each drug). Dose-response curves of each drug showing dose-dependent effects of each drug on ovarian cancer cells. **(A)** CTG assay of TOV21G cell line treated with Ceralasertib with Dose-Response Curve (0-5µM). **(B)** CTG assay of OVCAR3 cell line treated with Ceralasertib with Dose-Response Curve (0-5µM). **(C)** CTG assay of Kuramochi cell line treated with Ceralasertib with Dose-Response Curve (0-160µM). **(D)** CTG assay of A2780 cell line treated with Ceralasertib with Dose-Response Curve (0-40µM). **(E)** CTG assay of CAOV3 cell line treated with Ceralasertib with Dose-Response Curve (0-40µM). (**F)** CTG assay of SKOV3 cell line treated with Ceralasertib with Dose-Response Curve (0-40µM). **(G)** CTG assay of TOV21G cell line treated with ONC201 with Dose-Response Curve (0-40µM). **(H)** CTG assay of OVCAR3 cell line treated with ONC201 with Dose-Response Curve (0-40µM). **(I)** CTG assay of Kuramochi cell line treated with ONC201 with Dose-Response Curve (0-80µM). **(J)** CTG assay of A2780 cell line treated with ONC201 with Dose-Response Curve (0-60µM). **(K)** CTG assay of CAOV3 cell line treated with ONC201with Dose-Response Curve (0-60µM). **(L)** CTG assay of cell line treated with ONC201 with Dose-Response Curve (0-160µM).

**Table 1.** Histological type and known *TP53* status and ic50 values of Ceralasertib and ONC201 in ovarian cancer cell lines.

| **Cell line** | **Histological type** | ***TP53* status** | **Ceralasertib ic50 (μM)** | **Onc201 ic50 (μM)** |
| --- | --- | --- | --- | --- |
| TOV21G | Non serous | Wild type | 0.84 | 2.285 |
| OVCAR3 | High grade serous | mutated | 0.55 | 0.88 |
| KURAMOCHI | High grade serous | mutated | 54.75 | 2.31 |
| A2780 | Non serous | Wild type | 0.92 | 1.715 |
| CAOV3 | High grade serous | mutated | 0.55 | 1.67 |
| SKOV3 | Non serous | P53 null | 1.48 | 20.16 |

We did not observe more sensitivity toward the ATRi ceralasertib in p53-mutated tumor cells as compared to cell lines with wild-type p53. The Kuramochi high grade serous ovarian cancer cell line with mutated p53 had a higher IC50 value of ceralasertib compared to other cell lines.

### Synergistic cell killing effect at 72 hours from combination ONC201 and ceralasertib in ovarian cancer cell lines

We combined ONC201 and ATRi ceralasertib to assess the effect of combination on cell viability of the ovarian cancer cell lines. **Figure 2** shows combinational synergy with scores less than one using multiple doses of both drugs as calculated by Compusyn software.

**Figure 2.**
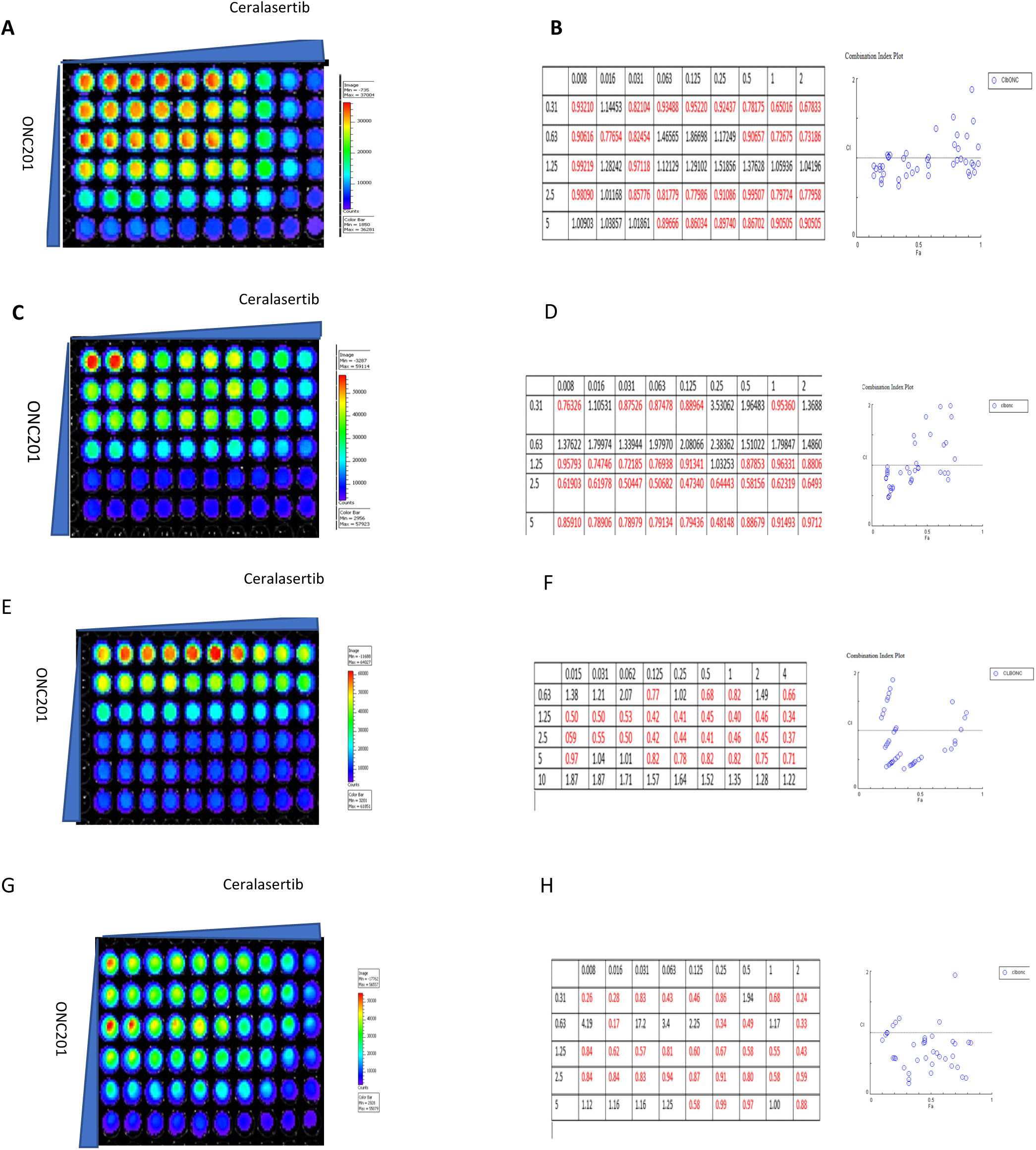
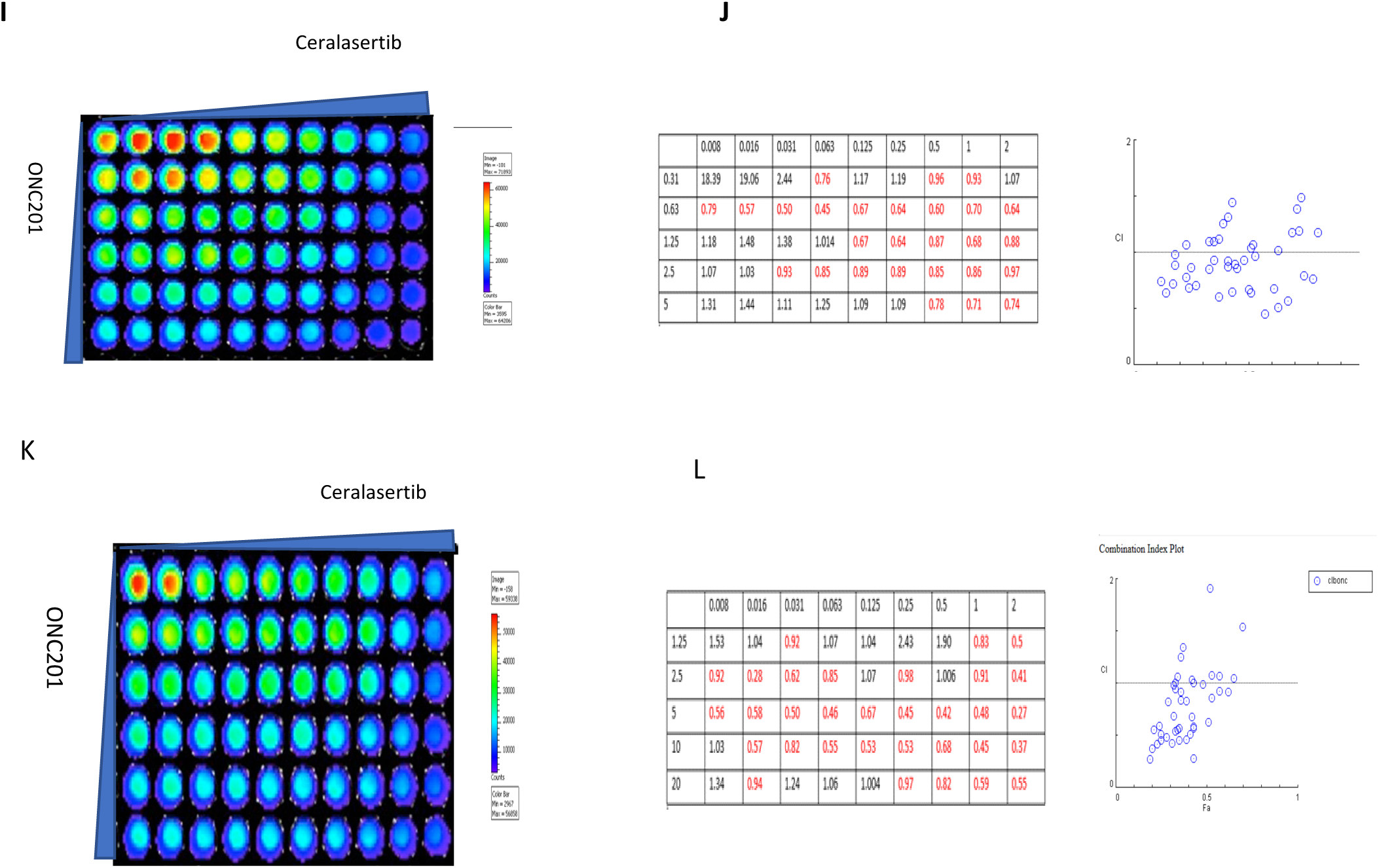
Ceralasertib and ONC201 synergistically reduce cell viability in ovarian cancer cells. TOV21G, OVCAR3, Kuramochi, A2780, CAOV3, SKOV3 cell lines were treated with increasing doses of Ceralasertib and ONC201 and cell titer Glo assay was performed after 72 hours of treatment as described in Methods. Synergy scores and combination index plot of each cell line calculated by Compusyn software are also shown for each cell line. (**A)** CTG assay of TOV21G cell line treated with increasing dosage of Ceralasertib and ONC201(Ceralasertib: 0-2 µM, ONC201: 0-5µM). **(B)** Synergy Score and combination index plots of TOV21G cell line treated with Ceralasertib-ONC201 (numbers <1.0 show synergy). **(C)** CTG assay of OVCAR3 cell line treated with increasing dosage of Ceralasertib and ONC201 (Ceralasertib: 0-2 µM, ONC201: 0-5µM). **(D)** Synergy Score and combination index plots of OVCAR3 cell line treated with Ceralasertib-ONC201 (numbers <1.0 show synergy). **(E)** CTG assay of Kuramochi cell line treated with increasing dosage of Ceralasertib and ONC201 (Ceralasertib: 0-4 µM, ONC201: 0-10µM). **(F)** Synergy Score and combination index plots of Kuramochi cell line treated with Ceralasertib-ONC201(numbers <1.0 show synergy). **(G)** CTG assay of A2780 cell line treated with increasing dosage of Ceralasertib and ONC201(Ceralasertib: 0-2 µM, ONC201: 0-5µM). **(H)** Synergy Score and combination index plots of A2780 cell line treated with Ceralasertib-ONC201 (numbers <1.0 show synergy). **(I)** CTG assay of CAOV3 cell line treated with increasing dosage of Ceralasertib and ONC201(Ceralasertib: 0-2 µM, ONC201: 0-5µM), **(J)** Synergy Score and combination index plots of CAOV3 cell line treated with Ceralasertib-ONC201 (numbers <1.0 show synergy). **(K)** CTG assay of SKOV3 cell line treated with increasing dosage of Ceralasertib and ONC201(Ceralasertib: 0-2 µM, ONC201: 0-20 µM). **(L)** Synergy Score and combination index plots of SKOV3 cell line treated with Ceralasertib-ONC201 (numbers <1.0 show synergy).

In the OVCAR3 cell line the highest synergy score was observed at 0.125 μM ceralasertib and 2.5 μM ONC201. In the TOV21G cell line the highest synergy score was observed at a concentration of 1 μM ceralasertib and 0.31 μM ONC201. In the Kuramochi cell line although ceralasertib IC50 was higher, smaller doses in combination with ONC201 increased killing of the cancer cells. The highest synergy score in this cell line was observed at 4 μM ceralasertib and 1.25 μM ONC201. In A2780 cells the ONC201 plus ceralasertib drug combination showed the best synergy with score of 0.24 at 2 μM ceralasertib and 0.31 μM ONC201. In the SKOV3 cell line the best synergy was observed at 2 μM ceralasertib and 5 μM ONC201. Finally, the CAOV3 cell line showed the best synergy at 0.063 μM ceralasertib and 0.63 μM ONC201.

### Ceralasertib and ONC201 combination enhanced synergistic apoptosis in ovarian cancer cells

We explored if the observed synergy between ONC201 and ATRi ceralasertib was caused by apoptosis in the ovarian cancer cell lines. As shown in **Figure 3**, combination of ONC201 and ceralasertib induced apoptosis as demonstrated by enhanced PARP cleavage. PARP cleavage occurred in the TOV21G cell line with different dosages of drugs at 48 or 72 hours after treatment in combination. PARP cleavage was also observed in the A2780 cell line after 48 hours of treatment in combination more than each drug alone. In the OVCAR3 cell line cPARP induction was observed as compared to control (**Figure 3**).

**Figure 3.**
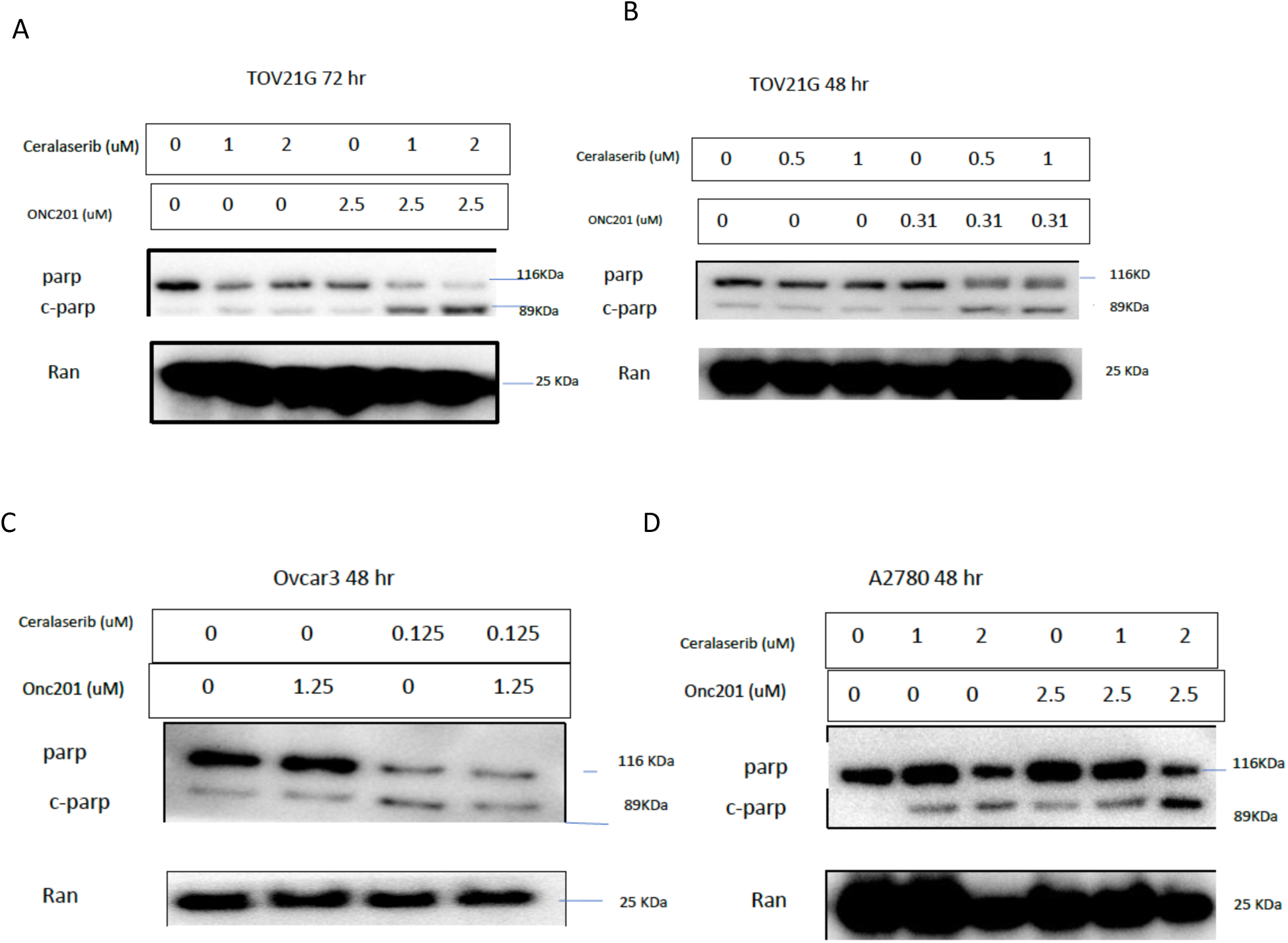
Ceralasertib and ONC201 combination induced PARP cleavage in ovarian cancer cells after 48 or 72 hours of treatment with different doses of drugs. **(A)** TOV21G cells treated 72 hr with Ceralasertib, ONC201 and combination. **(B)** TOV21G cells treated 48 hr with Ceralasertib, ONC201 and combination. **(C)** OVCAR3 cells treated 48 hr with Ceralasertib, ONC201 and combination. **(D)** A2780 cells treated 48 hr with Ceralasertib, ONC201 and combination

### Ceralasertib and ONC201 combination causes synergistic cell death in ovarian cancer cells associated with down regulation of anti-apoptotic Proteins

Bcl-XL as an anti-apoptotic protein was reduced in expression with combination of ONC201 and ceralasertib in all cell lines tested. Both ONC201 and ATRi could reduce Bcl-XL as single agents as shown in previous articles (21,22). Our western blot analysis showed the reduction of this marker with combination rather than each drug alone (**Figure 4**). Xiap as an X-linked inhibitor of apoptosis was decreased with single drug and combination. Xiap has been shown to be an important factor in cell survival of ovarian cancer leading to chemoresistance. The effect of ONC201 on Xiap was shown previously in pancreatic cancer (23, 24, **Figure 4**). cIAP-1, another inhibitor of apoptosis, was decreased with ONC201 and combination of ONC201 and ceralasertib. The effect of ONC201 in decreasing cIAP-1 was also described previously in pancreatic cancer cells (25, **Figure 3D**).

**Figure 4.**
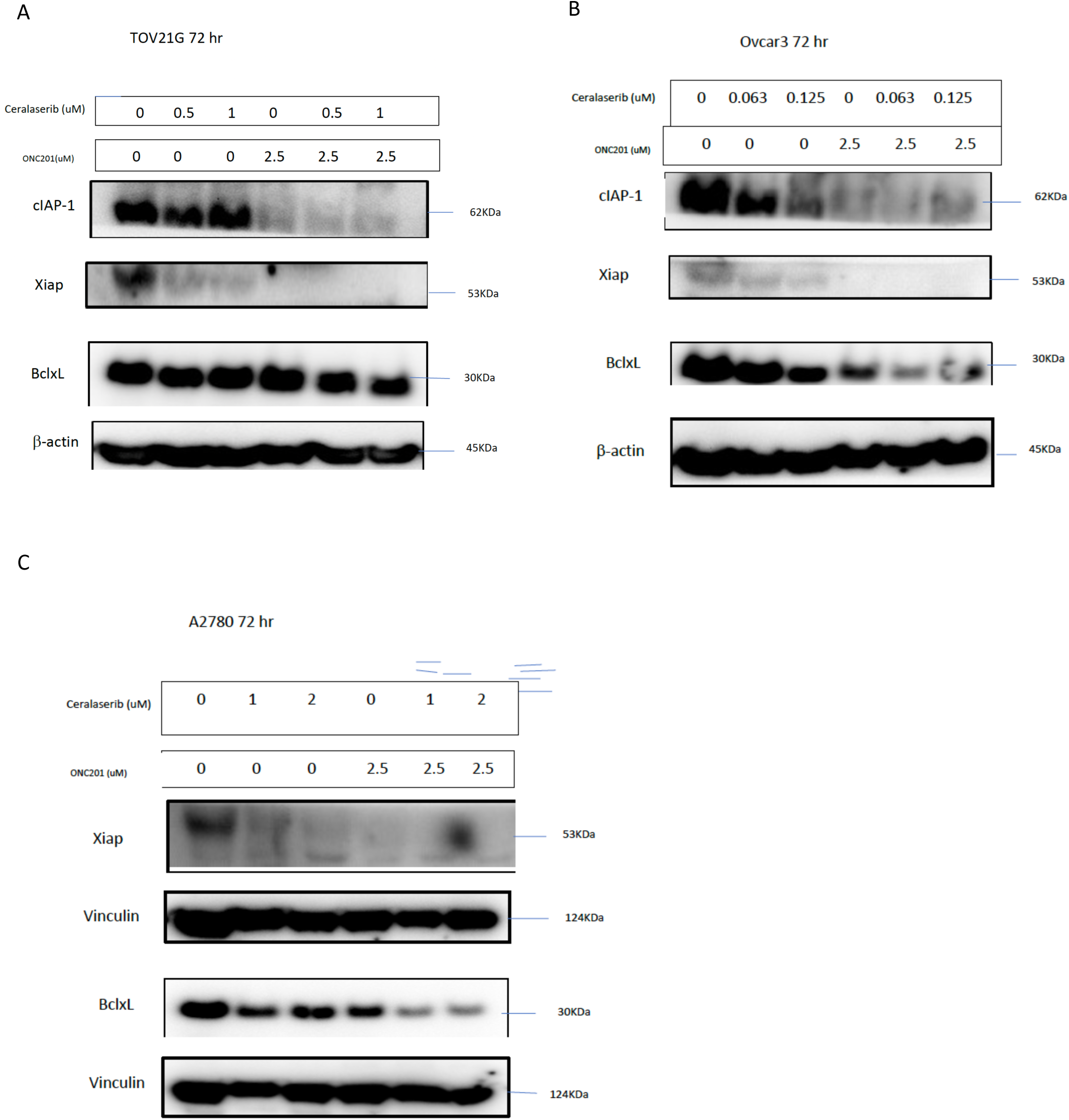
Ovarian cancer cell lines TOV21G, OVCAR3 and A2780 were treated with ATR inhibitor Ceralasertib and ONC201. **(A)** Tov21G cells treated with Ceralasertib, ONC201 and Combination for 72 hours**. (B)** OVCAR3 cell line was treated with Ceralasertib, ONC201 and combination for 72 hours. (C) A2780 cell line was treated with Ceralasertib, ONC201 and combination for 72 hours. Expression of BclxL as an anti-apoptotic protein was decreased with combination treatment after 72 hours. Xiap, a marker of cell survival was decreased with single treatments and combination in TOV21G, A2780 and OVCAR3 cell lines. cIAP-1 another marker of cell survival was decreased in TOV21G, OVCAR3 cell lines mostly with ONC201 single treatment. Each Membrane was probed once with loading control, some then were stripped and examined for other proteins (this didn’t happen more than once).

### Ceralasertib and ONC201 combination reduces p-Akt

We explored the effect of combination therapy of ONC201 and ceralasertib on a p-Akt. As mentioned earlier in the DDR, proteins like ATR and PARP can cause activation of Akt and the Akt pathway in turn can be a mechanism of resistance to DDR inhibitor therapy (18,19). ONC201 also can result in inactivation of the Akt signaling pathway,

Our investigation showed that combination of ONC201 and ceralasertib resulted in p-Akt reduction in the OVCAR3 and Kuramochi cell lines after 72 hours. In the TOV21G and A2780 cell lines, reduction in p-Akt was not observed at the dosage used after 72 hours of treatment (**Figure 5**).

**Figure 5.**
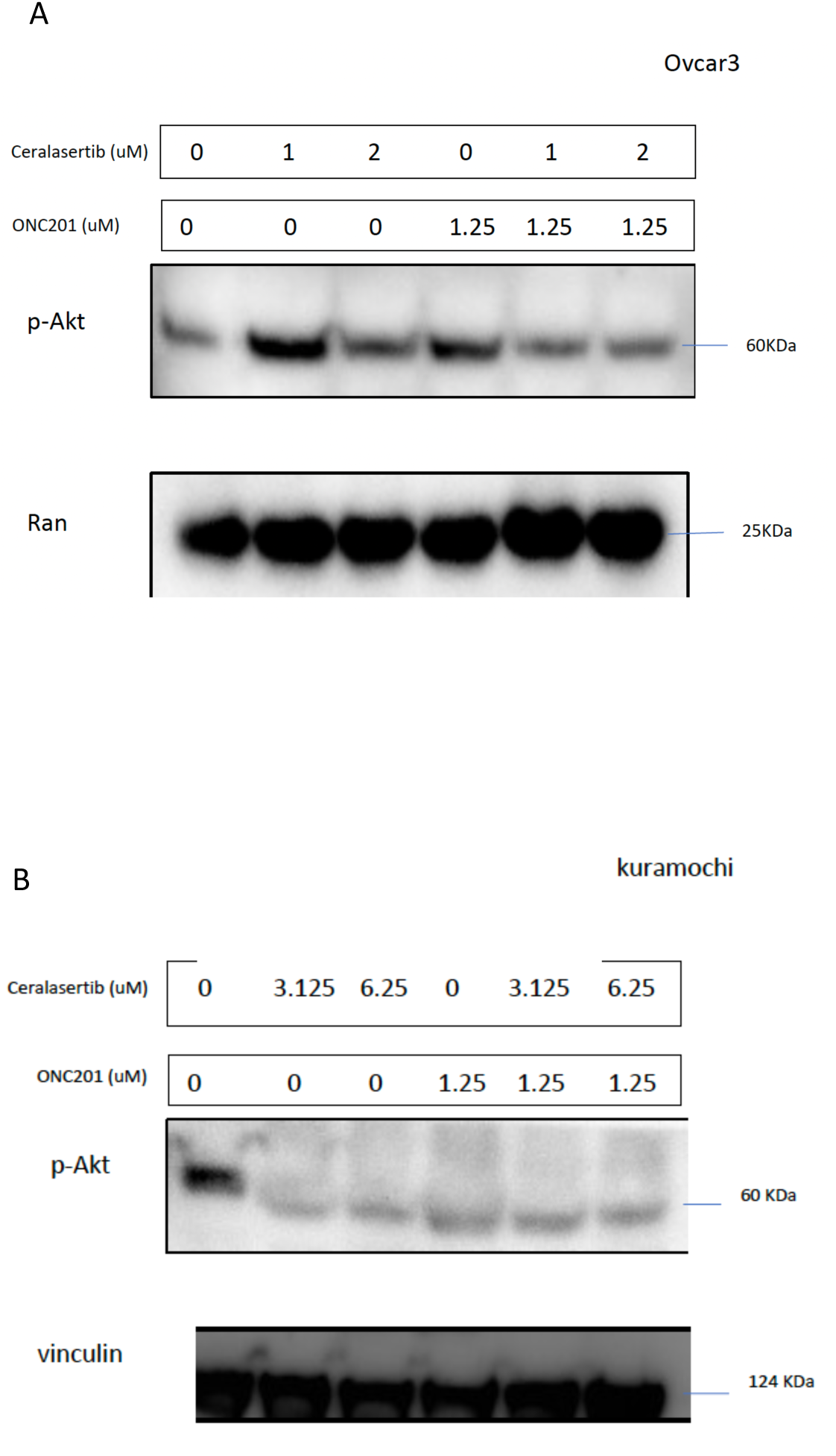
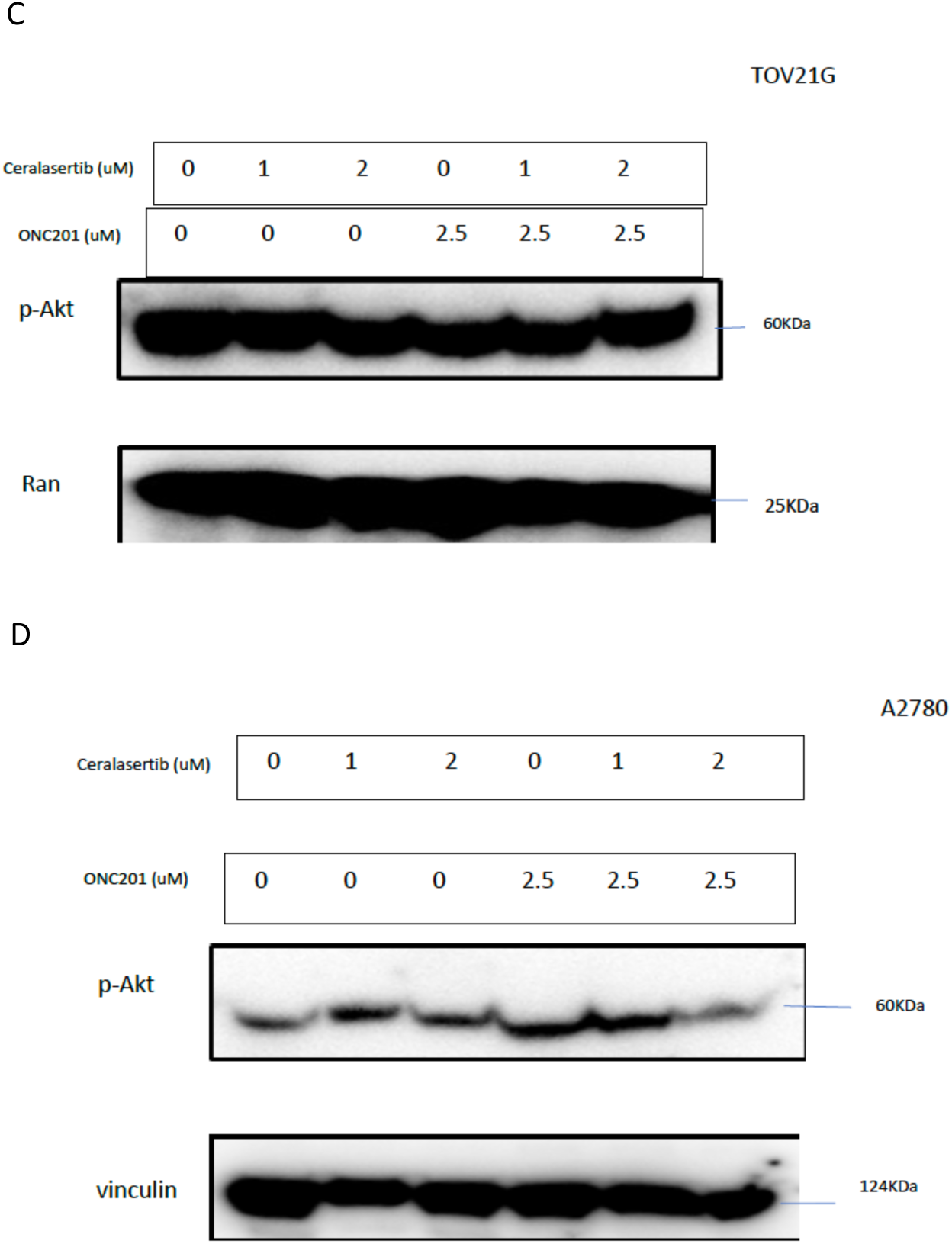
Combination of ONC201 and ATR inhibitor Ceralasertib decreased p-Akt expression in a heterogeneous manner among treated cells. **(A)** OVCAR3 cell line, treated with Ceralasertib, ONC201 and Combination. In ovcar3 cell line, proteins p-Akt and Ran are from the same membrane. **(B)** Kuramochi cell line, treated with Ceralasertib, ONC201 and combination. In Kuramochi cell line, proteins p-Akt and Vinculin are from the same membrane. **(C)** TOV21G cell line, treated with Ceralasertib, ONC201 and combination. In TOV21G cell line, proteins p-Akt and Ran and cleaved PARP (Figure 3) are from the same membrane. **(D)** A2780 cell line, treated with Ceralasertib, ONC201 and combination. In A2780 cell line, proteins p-Akt, vinculin, bclxl (Figure 4) are from same membrane.

### ONC201 caused increase in DR5 and ATF-4 expression as soon as 24 hours after treatment

Treatment with ONC201 previously showed an increase in markers of integrated response like ATF-4 even sooner than 24 hour of treatment and ATF-4 expression mediates an increase in DR5 expression through CHOP.

TOV21G (p53 wild-type cell line) was treated with ATRi ceralasertib and ONC201 individually and in combination. ONC201 increased in ATF-4 and DR5 expression as a single treatment and in combination treatment after 24 hours of treatment and this increase was greater in the combination group (**Figure 6**).

**Figure 6.**
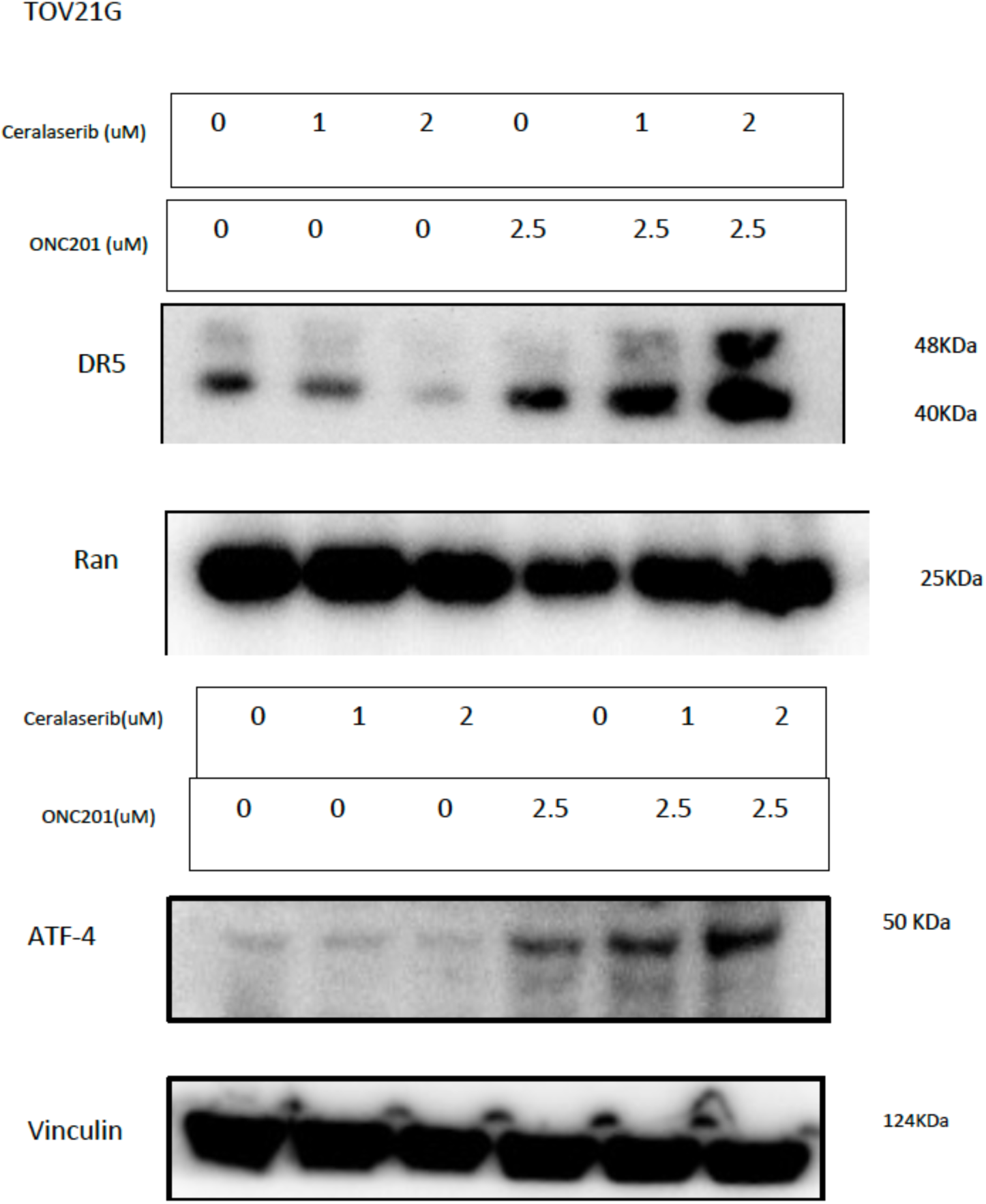
**Increase in expression of DR5 and ATF-4 in ONC201 and combination treatment at 24 hours in TOV21G cell line.**

### CLPX expression was reduced with ONC201 monotherapy and with combination of ONC201 and ceralasertib as early as 24 hours

Imipridones previously showed a strong effect on CLPX. CLpP and its companion CLPX form a complex named CLpXP that degrades misfolded Proteins (26). Knowing the effect of imipridones on CLPX can occur as early as 6 hours (27), we examined this effect in the TOV21G and A2780 cell lines after 24 and 48 hours of treatment. ONC201 decrease CLPX levels at 24 hours of treatment in the drug only and combination of ONC201 and ceralasertib. The effect was observed at 48 hours as well. ATRi ceralasertib also decreased CLPX levels after 48 hours of treatment in both cell lines in a dose dependent manner (**Figure 7**).

**Figure 7.**
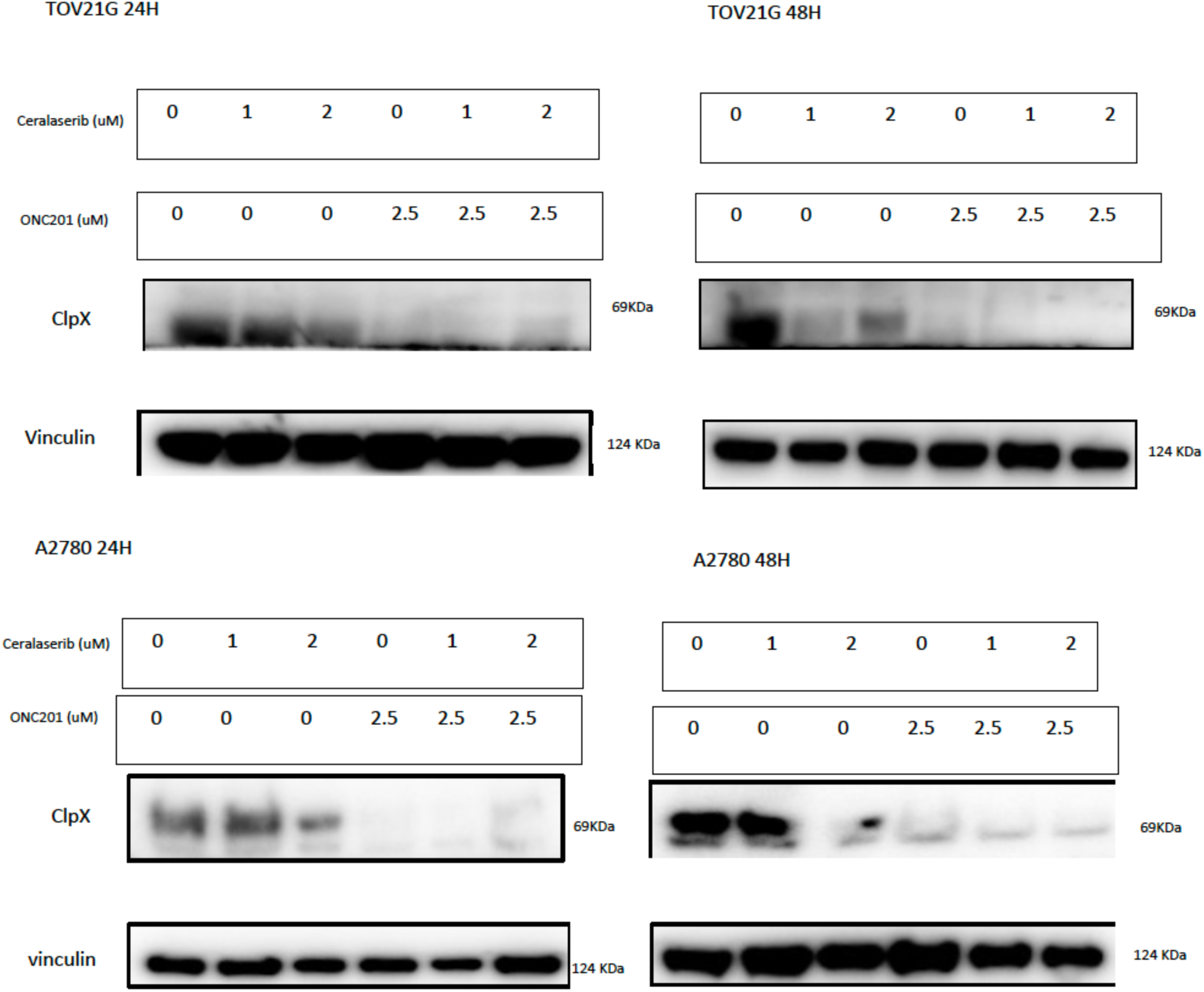
The expression of ClpX decreases following ONC201 and combination treatments. ATR inhibitor Ceralasertib also decreases ClpX expression in a dose-dependent manner.

### Cytokine profiling showed decreased in markers of inflammation and tumor progression in ONC201 and ceralasertib treated ovarian cancer cell lines

The Fold-change value of each treatment compared to NO-treatment were calculated and a Heat map was drawn for each cell line and time point (**Figure 8A**). Cytokine analysis in the TOV21G cell line after 48 hours of treatment with ONC201, ceralasertib and combination showed decrease in CCL2 (**Figure 8B**). In general, CCL2 is a marker of tumor growth and metastasis and leads to poor prognosis in cancer patients because of an immunosuppressive tumor environment (28).

**Figure 8.**
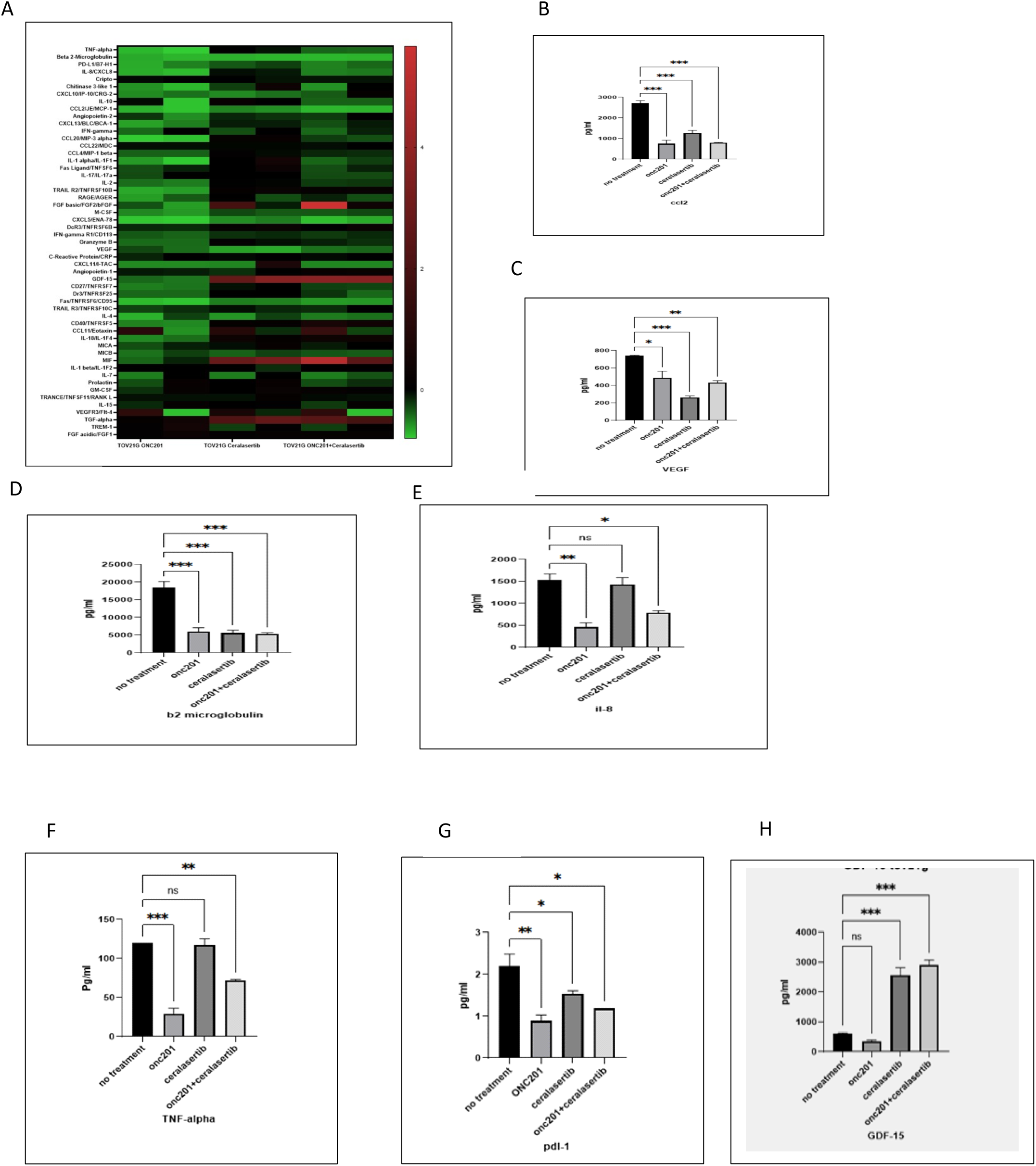
Cytokine alterations in ONC201, Ceralasertib and combination treated cells. **(A)** Heat map of fold-change analysis of cytokines produced in tumor microenvironment of TOV21G cell line treated with ONC201, Ceralasertib and Combination after 48 hours. There are biological replicates of each condition. **(B)** CCL2 was decreased as a marker of tumor progression and inflammation after treatment with ONC201, Ceralasertib and combination in TOV21G cell line after 48 hours of treatment. **(C)** VEGF was decreased as a marker of tumor progression and inflammation after treatment with ONC201, Ceralasertib and combination in TOV21G cell line after 48 hours of treatment. **(D)** β2-mIcroglobulin was decreased as a marker of tumor progression and inflammation after treatment with ONC201, Ceralasertib and combination in TOV21G cell line after 48 hours of treatment. **(E)** IL-8 as other marker of tumor progression in ovarian cancer was decreased after ONC201 and Combination treatment in TOV21G cell line after 48 hours of treatment. **(F)** TNF-alpha which can lead to other chemokine productions that can cause tumor progression by ovarian cancer cells is decreased with ONC201 treatment and combination in TOV21G cell line after 48 hours of treatment. **(G)** PDL-1 as one of the regulators of cancer immune response was decreased after treatment with ONC201, Ceralasertib and combination in TOV21G cell line after 48 hours of treatment. **(H)** GDF-15 as a marker of immune escape and a predictor of chemotherapy response was increased after treatment with ATR inhibitor ceralasertib and as a result in combination of Ceralasertib and ONC201.

VEGF levels also went down after all 3 conditions (**Figure 8C**). VEGF is an angiogenesis cytokine that contributes to tumor progression and growth in ovarian cancer specifically; VEGF is a contributing factor to tumor progression and metastasis and peritoneal spreading of ovarian tumors (29). Beta2 microglobulin also decreased after ONC201, ceralasertib or combination therapy (Figure 8D). Beta2 microglobulin is another marker contributing to tumor progression and potential therapeutic biomarker of cancer (30). Beta2 microglobulin was more expressed in ovarian tumors compared to normal cells and lower beta2 microglobulin caused less tumor progression and migration (31).

IL-8 and TNF-alpha were decreased after ONC201 treatment alone or combination of ONC201 and ceralasertib treatment (**Figure 8E, F**) as compared to control and ceralasertib alone. IL-8 was demonstrated to increase ovarian tumor progression and invasion with a higher level of IL-8 in ovarian cancer patients leading to poor Prognosis (32,33). TNF-alpha made by ovarian cancer cells can lead to other chemokine production such as CCL2, IL-6 and VEGF resulting in peritoneal invasion of tumor (34).

PD-L1 was also decreased with single treatments and combination of ONC201 and ceralasertib. As shown before ATRi can downregulate PD-L1 and caused enhanced T-cell mediated killing of cancer cells (35). This result also shows the effect of ONC201 and combination of ONC201 and ceralasertib on downregulating PD-L1 and possible positive effect of these drugs and combination on immune mediated killing of cancer cells because of the important role of PD-L1/PD-1 axis in immune escape of cancer cells (36, **Figure 8G**).

GDF-15 was increased after ceralasertib treatment alone or after combination of ONC201 and ceralasertib treatment in the p53 wild-type TOV21G cell line (**Figure 8H**). GDF-15, a p53 target gene (37), is a member of Transforming growth factor beta (TGF-β) super family (38,39). GDF-15 was shown to predict chemotherapy response in ovarian cancer with the level being higher in patients with chemoresistance (40). GDF-15 was also shown to help tumor growth and immune escape (41).

In the OVCAR3 cell line, cytokine profiling was performed after 24h of treatment (**Figure 9A**) and showed that Angiopoeitin-2 was meaningfully decreased in OVCAR3 cells after ceralasertib alone or combination of ONC201 and ceralasertib treatment (**Figure 9B**). Angiopoietins are one of the chemo markers of tumor progression. In ovarian cancer especially it was shown that they can lead to tumor progression, metastasis and peritoneal involvement (42,43). IL-6 as a negative marker causing progression and therapy resistance was also increased after ONC201 treatment alone and as a result in combination of ONC201 and ceralasertib treatment (**Figure 9C**, 44).

**Figure 9.**
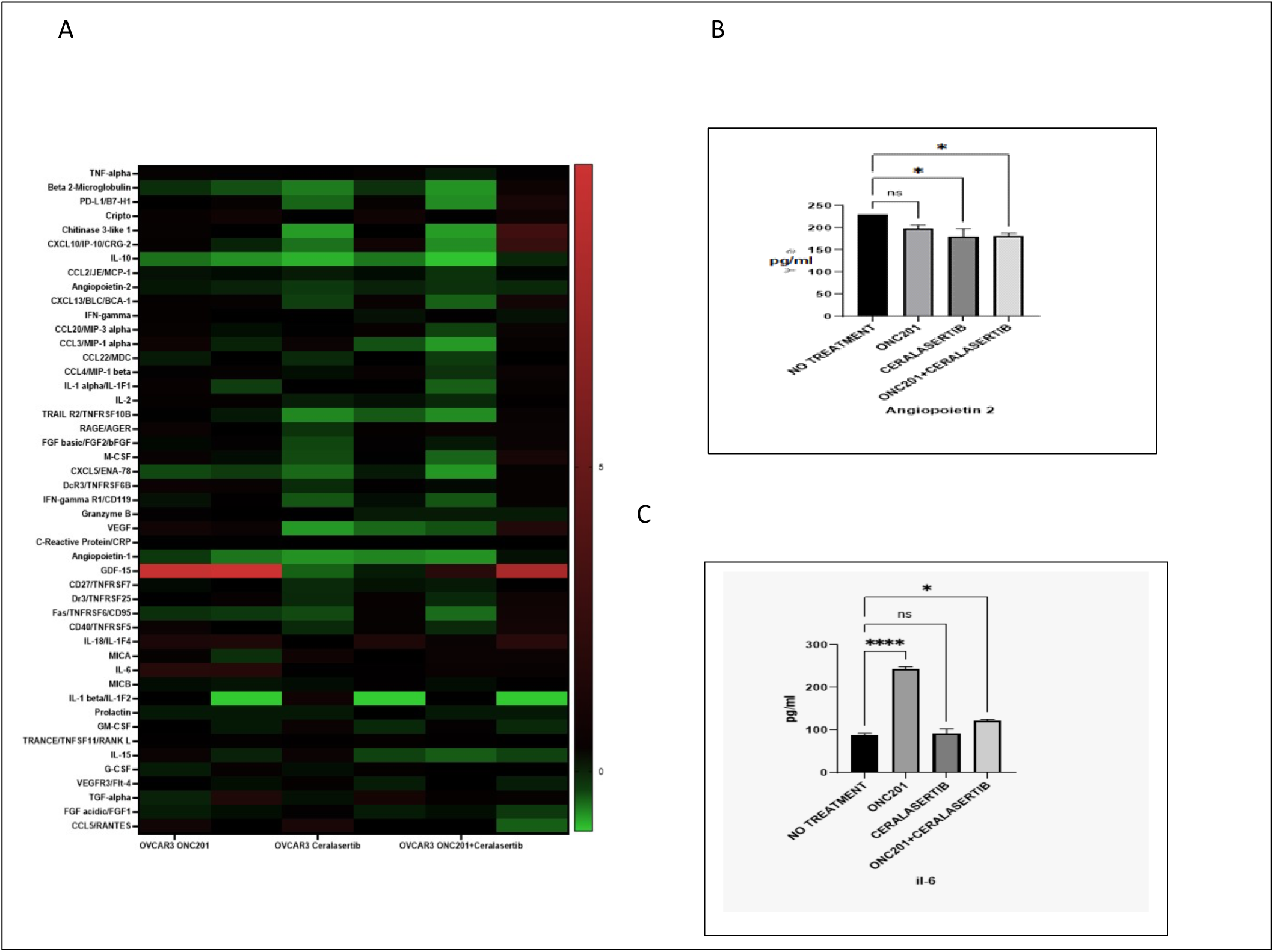
Cytokine changes in ovcar3-treated cells. **(A)** Heat map of fold-change analysis of cytokines produced in tumor microenvironment of ovcar3 cell line treated with ONC201, Ceralasertib and Combination after 24 hours. **(B)** Angiopoietin 2 as a marker of tumor progressions was decreased at ceralasertib and combination in ovcar3 cell line after 24 hours of treatment. **(C)** IL6 another marker of tumor progression as a negative marker was increased with ONC201 and as a result in combination in ovcar3 cell line after 24 hours of treatment.

## Discussion

Treatment of ovarian cancer faces challenges because of tumor recurrence and resistance to initial treatment. New treatment options need to be developed to enhance treatment efficacy and response.

Ovarian cancer and especially high grade serous ovarian cancer suffer from dysfunctional p53. Almost all high-grade serous type ovarian cancers have a dysfunctional p53 pathway which makes this cancer a good target for ATRi (45,46). ATRi have shown different results in regard to p53 status. *TP53* mutation was not associated with higher sensitivity to ATRi VE-821 but HGSOC cells were more sensitive to the compound (47).

ATRi are now widely tested in clinical trials mostly in combination with other drugs and modalities like PARP inhibitors, chemo- and radio-therapy (11, 13). One of the challenges of treatment with ATRi is adverse hematological side effects observed in combination with chemotherapeutics (48). Thus it is important to have new tolerable combinations to enhance drug efficacy and reduce the side effects.

ONC201, a TRAIL-Inducing Compound (TIC10), has shown efficacy in ovarian cancer cells and mouse models (21). ONC201 is a well-tolerated compound as observed in clinical trials for solid tumors.

In this study we tested the combination of ATRi ceralasertib and ONC201 against 6 ovarian cancer cell lines 3 high grade serous and 3 non-serous ovarian cancer cell lines. Two of the cell lines were p53 wild-type (TOV21G, A2780) and 3 harbored mutated p53 (OVCAR3, KURAMOCHI, CAOV3) and one was p53-null (SKOV3). both ceralasertib and ONC201 were effective as monotherapy against ovarian cancer cell lines. Cell lines with wild-type p53 were as sensitive to ATRi ceralasertib as those with p53 mutations. The IC50 ranged between 0.55-54.75 μM. All examined cell lines regardless of histological type and p53 status were sensitive to ATRi ceralasertib. Only the KURAMOCHI cell line, a high grade serous ovarian cancer cell line with p53 mutation showed a much higher IC50 value as compared to other cell lines (**Table 1** and **Figure 1**). Viability assays showed that the two drug ceralasertib and ONC201 combination caused synergy among ovarian cancer cells after 72 hours at multiple dosages (**Figure 2**). Western blot analysis showed that the ceralasertib and ONC201 combination caused cPARP induction (**Figure 3**), reduced Bcl-XL (**Figure 4**), and Xiap reduction (**Figure 4**) consistent with induction of apoptosis. cIAP-1 was decreased mostly as with ONC201 treatment in the TOV21G and OVCAR3 cell lines (**Figure 4**).

Our investigation of a common pathway between ceralasertib and ONC201 and the reason behind their synergy suggested that the combination can decrease p-Akt (**Figure 5**). Reduced p-Akt causes inactivation of the Akt signaling pathway which has an important role in ovarian cancer development and resistance to therapy, especially with DDRi’s. It is worth noting that the expression of apoptotic markers and p-Akt was heterogeneous among cancer cells.

Looking into the effect of ceralasertib and ONC201 combination on the integrated stress response (ISR) we examined the effect of combination on ATF-4 and subsequently on TRAIL receptor DR5 expression as early as 24 hours in the TOV21G cell line. ONC201 increased ATF-4 expression in ONC201-treated cells and ceralasertib and ONC201 combination showed slightly more increase in ATF-4 (Figure 6).

Our investigation on effect of ceralasertib and ONC201 treatment on CLPX expression demonstrated decrease in CLPX level at 24 hours in ONC201-treated and combination group. ATRi ceralasertib also contributed in CLPX reduction after 48 hours of treatment. The decrease in CLPX level after ceralasertib occurred after 48 hours in a dose-dependent manner (**Figure 7**). CLPX is a member of CLPXP complex which has a role in degradation of misfolded proteins in mitochondria (26). By decreasing CLPX levels the ceralasertib and ONC201 combination disrupts mitochondrial function and can cause growth arrest. Studies have suggested the importance of mitochondrial function in ovarian cancer. Mitochondrial function can be elevated in some ovarian cancer cells making them possibly more sensitive to treatments that disrupt its function (49).

Our studies into ceralasertib and ONC201 effects on the tumor microenvironment by cytokine profiling showed that both drugs individually and in combination cause decreases in markers of tumor progression and inflammation like CCL2, beta2 microglobulin, VEGF (**Figures 8B, 8C, 8D**) (28, 29, 30, 31).

Markers including IL-8, and TNF-alpha that can lead to production of other inflammatory cytokines were reduced with ONC201 and the ceralasertib and ONC201 combination (**Figures 8E, 8F**) (32, 33, 34).

PD-L1, an important marker in tumor-immune response, was reduced with ceralasertib and ONC201 monotherapy or combination of ceralasertib and ONC201 (**Figure 8G**). PD-L1 has an important role in immune response of cancer cells (35).

Interestingly we observed an increase in GDF-15 with ATRi treatment or ceralasertib and ONC201 combination treatment in the p53 wild-type TOV21G cell line (**Figure 8H**). GDF-15 is a p53 target gene which has an important role in ovarian cancer progression and immune escape (37, 38, 39, 40, 41). This data suggests the possibility of combining an ATRi and GDF-15 inhibitor to increase treatment efficacy and immune response in ovarian cancer.

Overall, our data demonstrates a synergy between ATRi ceralasertib and ONC201 that can be translated into patients. To our knowledge, this is the first study showing the effectiveness of ceralasertib and ONC201 combination in ovarian cancer *in vitro*. The combination of ceralasertib and ONC201 needs further studies to confirm the effect of treatment *in vivo* and in clinical trials.

## Acknowledgements

W.S.E-D. is an American Cancer Society Research Professor and is supported by the Mencoff Family University Professorship at Brown University. This work was supported by an NIH grant (CA173453) to W.S.E-D. and by funding from Chimerix, Inc. This work was presented in part at the 2023 meeting of the American Association for Cancer Research.

## Disclosure of conflict of interest

W.S.E-D. founded Oncoceutics, Inc. in 2004. Oncoeutics later licensed TIC10/ONC201 originally discovered in his lab in 2007. Oncoceutics was acquired by Chimerix in 2021. Chimerix was subsequently acquired by Jazz Pharmaceuticals in 2025 and took ONC201 to FDA approval as dordaviprone. W.S.E-D. is a founder of p53-Therapeutics, Inc. in 2013, Inc., a biotech company focused on developing novel small molecule anti-cancer therapies targeting mutant p53 protein. He founded SMURF-Therapeutics, Inc. in 2021, a biotech company focused on developing therapeutics targeting HIF1-alpha, including a micro-RNA that targets CDK4/6 to destabilize HIF. Dr. El-Deiry has disclosed his entrepreneurial relationships and potential conflicts of interest to his academic institution/employer and is fully compliant with institutional and NIH policy that is managing this potential conflict of interest.

